# Peripheral infrastructure vectors and an extended set of plant parts for the modular cloning system

**DOI:** 10.1101/237768

**Authors:** Johannes Gantner, Theresa Ilse, Jana Ordon, Carola Kretschmer, Ramona Gruetzner, Christian Löfke, Yasin Dagdas, Katharina Bürstenbinder, Sylvestre Marillonnet, Johannes Stuttmann

## Abstract

Standardized DNA assembly strategies facilitate the generation of multigene constructs from collections of building blocks in plant synthetic biology. A common syntax for hierarchical DNA assembly following the Golden Gate principle employing Type IIs restriction endonucleases was recently developed, and underlies the Modular Cloning and GoldenBraid systems. In these systems, transcriptional units and/or multigene constructs are assembled from libraries of standardized building blocks, also referred to as phytobricks, in several hierarchical levels and by iterative Golden Gate reactions. This combinatorial assembly strategy meets the increasingly complex demands in biotechnology and bioengineering, and also represents a cost-efficient and versatile alternative to previous molecular cloning techniques. For Modular Cloning, a collection of commonly used Plant Parts was previously released together with the Modular Cloning toolkit itself, which largely facilitated the adoption of this cloning system in the research community. Here, a collection of approximately 80 additional phytobricks is provided. These phytobricks comprise e.g. modules for inducible expression systems, different promoters or epitope tags, which will increase the versatility of Modular Cloning-based DNA assemblies. Furthermore, first instances of a “peripheral infrastructure” around Modular Cloning are presented: While available toolkits are designed for the assembly of plant transformation constructs, vectors were created to also use coding sequence-containing phytobricks directly in yeast two hybrid interaction or bacterial infection assays. Additionally, DNA modules and assembly strategies for connecting Modular Cloning with Gateway Cloning are presented, which may serve as an interface between available resources and newly adopted hierarchical assembly strategies. The presented material will be provided as a toolkit to the plant research community and will further enhance the usefulness and versatility of Modular Cloning.

## Introduction

Molecular cloning belongs to the unbeloved, yet inevitable everyday tasks of many wet lab molecular biologists. In the past two decades, most labs either relied on classical ligation of restriction fragments or PCR products into a vector of interest, or used the gateway system based on the recombination reactions taking place during the integration and excision of the genome of phage lambda in bacterial infection (Katzen, 2007). Gateway cloning proved to be extraordinarily efficient for regular cloning, but also for high-throughput applications such as library generation, and tremendously reduced molecular cloning workloads. One striking advantage of gateway cloning is that only the initial creation of entry clones represents a critical step. Subsequently, inserts may be mobilized from entry clones into a wide array of destination vectors by basically failsafe, highly efficient and unified recombination reactions. This is also facilitated by the availability of destination vectors for virtually any biological system and experimental setup (e.g. Earley et al., 2006; Nakagawa et al., 2007). However, Gateway cloning is relatively costly, as it relies on use of the proprietary BP/LR enzyme blends, and also represents a rather binary approach to molecular cloning where a single insert is mobilized into a new sequence context. Theoretically, this was overcome by the invention of multisite gateway systems (Cheo et al., 2004). The current MultiSite Gateway™ Pro technology allows combination of up to four DNA fragments in any (attR1/R2 site-containing) destination plasmid. Multisite gateway was also combined with other cloning techniques in Golden GATEway cloning for further flexibility and generation of multigene constructs (Kirchmaier et al., 2013). However, the multisite gateway technology found only limited use in the scientific community, as novel DNA assembly strategies concomitantly emerged. Most popular strategies for combinatorial DNA assembly now rely on Gibson assembly or Golden Gate cloning.

In Gibson assembly (Gibson et al., 2009), linear DNA fragments sharing identical sequence stretches of e.g. 20 - 30 base pairs at their ends are stitched together in single tube reaction. First, 5’ ends of DNA fragments are chewed back to create single strand 3’ overhangs by an exonuclease. By the identical sequence ends, complementary fragments anneal. The annealed fragments are then covalently fused by a polymerase filling up gaps and a ligase removing nicks. Thus, fusion sites between fragments in Gibson assembly are scarless, no particular sequence motifs such as restriction sites are required, and large sequences up to several hundred kilobases can be assembled (Gibson et al., 2009). However, Gibson assembly relies on the engineering of identical ends on sequence fragments, and does therefore not provide a theoretical framework for re-utilization of DNA modules in multiple and diverse DNA assemblies. This was recently achieved by the invention of hierarchical DNA assembly strategies based on Golden Gate cloning (Engler et al., 2008). Three major standards, GreenGate (Lampropoulos et al., 2013), GoldenBraid (Sarrion-Perdigones et al., 2011) and Modular Cloning (Weber et al., 2011) were developed in parallel and are now used in the plant research community. All systems utilize the same basic principles: Standardized four base pair (bp) overhangs (generated by Type IIs restriction endonucleases) are defined as fusion sites between building blocks of transcriptional units, such as promoters, untranslated regions, signal peptides, coding sequences, or terminators. Building blocks are cloned as Level 0 modules, which are flanked by these four bp overhangs and recognition sites for a given Type IIs endonuclease. These units are also referred to as phytobricks. In a second hierarchical level (Level 1), phytobricks are assembled into transcriptional units by highly efficient Golden Gate cloning. Multigene constructs are assembled with another Golden Gate reaction and a yet further hierarchical level (Level 2 or Level M). Each system has its individual advantages and constraints. In GreenGate cloning, only the Type IIs enzyme *BsaI* is used. Thus, internal recognition sites of solely this enzyme have to be removed from building blocks during “sequence domestication” (elimination of internal Type IIs enzyme recognition sites). However, Level 1 vectors of the GreenGate system are not plant transformation vectors, and an additional assembly step in a Level 2 “destination” vector is thus required prior to functional verification. Furthermore, multigene construct assembly is carried out by iterative rounds where additional transcriptional units (Level 1) are added to an existing (Level 2) construct. Both GoldenBraid and Modular Cloning rely on cloning steps of different hierarchical levels being carried out by iterative use of two different Type IIs enzymes – BsaI and *BpiI* in Modular Cloning, and BsaI and *BsmBI* for GoldenBraid. This obviously increases the requirement for sequence modifications during domestication, but streamlines cloning procedures and increases flexibility. The developers of GoldenBraid and Modular Cloning also agreed on a common set of fusion sites between building blocks, a common “grammar” or syntax, making these systems at least partially compatible (Sarrion-Perdigones et al., 2013; Engler et al., 2014). The main difference between systems consists in the approach for multigene construct assembly: Being an iterative process in GoldenBraid, up to six Level 1 modules may be assembled in a Level 2 construct in a single step by Modular Cloning. This strategy facilitates and/or accelerates the assembly of multigene constructs, but comes at the expense of a more complex nomenclature and vector toolkit. GoldenBraid developers also provide online databases and software suites for end-users (Vazquez-Vilar et al., 2017), and similar tools are yet unavailable for Modular Cloning. A number of research laboratories recently agreed on the use of the common molecular syntax underlying both Modular Cloning and GoldenBraid to foster re-utilization and sharing of DNA modules for bioengineering (Patron et al., 2015).

The previously released Modular Cloning Toolkit provides DNA modules facilitating domestication of novel sequences and assembly of multigene constructs following the Modular Cloning standard (Engler et al., 2014). A simultaneously released collection of Plant Parts contains 95 modules coding for commonly used promoters, transcriptional terminators, epitope tags and reporter genes (Engler et al., 2014). In conjunction, these toolkits allow for a jump start into hierarchical DNA assembly for end users. In principle, the cloning of *your favorite gene (YFG)* in the Modular Cloning format (as a CDS1 or CDS1ns module) will be sufficient for assembly of *YFG* together with different Plant Parts in various simple or complex plant transformation constructs. However, a peripheral infrastructure which allows re-using the Modular Cloning *YFG* modules (CDS1 or CDS1ns Level 0 modules) in other experimental setups, such as e.g. bacterial or yeast expression, and also an interface to Gateway cloning strategies, were so far missing. Here, we present molecular tools for connecting the Modular Cloning system with Gateway cloning, either for toggling between Modular Cloning and Gateway cloning, the cost-efficient generation of Gateway entry clones, or simple, hierarchical assembly of Gateway destination vectors. Furthermore, vectors were developed for re-utilization of Modular Cloning *YFG* modules for yeast two hybrid assays or bacterial translocation into plant cells. Finally, an extended collection of Plant Parts consisting of ~ 80 Level 0 modules, or phytobricks, is provided for the sake of efficient bioengineering through shared resources.

## Results and Discussion

### Golden Gate cloning vectors for cost-efficient generation of gateway entry clones or shuttling from Modular Cloning to Gateway Cloning

Gateway entry clones can be generated by TOPO cloning, BP recombination reaction or classical ligation of restriction fragments (Katzen, 2007). TOPO cloning and BP recombination are comparatively expensive (~ 35 and 13 €/reaction list price, respectively). Classical ligation into e.g. pENTR11-like vectors requires individualized cloning strategies for different amplicons, and may also result in different linkers occurring in final fusion proteins depending on restriction sites used. We designed recipient vectors (pJOG130 and pJOG131) which are comparable to pENTR11, but sequences are inserted by Golden Gate cloning via *Bsa*I to generate entry clones (Fig. 1). Vectors are based on a common backbone (Kanamycin resistance, cloning cassette flanked by M13fwd and M13rev priming sites), but differ in overhangs generated by *Bsa*I digestion. pJOG130 uses overhangs of CDS1 modules of the Modular Cloning system (Fig. 1a; Weber et al., 2011; Engler et al., 2014). It can thus be used to convert Level 0 CDS1 modules of the Modular Cloning system to Gateway entry clones by simple *Bsa*I cut/ligation, or for cloning of PCR products flanked by adaptors to generate respective overhangs (Fig. S1a). pJ0G131 incorporates overhangs of CDS1ns modules of the Modular Cloning system. Correspondingly, it can be used to shuttle CDS1ns modules to the Gateway cloning system, or to clone PCR products carrying suitable adaptors (Fig. S1b). While both vectors may theoretically be used to clone PCR products containing or not a STOP codon, we intended to use pJOG130 for cloning of coding sequences with, and pJ0G131 for sequences without a STOP codon to follow a common nomenclature. Fusion sites resulting from recombination of inserts from pJOG130 and 131 into Gateway expression vectors are depicted in Fig. 1 c and d. *Att* site-flanking *AscI* and *NotI* sites present in most entry plasmids are maintained, and fusion sites from Golden Gate cloning translate into serine or alanine residues commonly employed as linkers.

**Figure 1:**
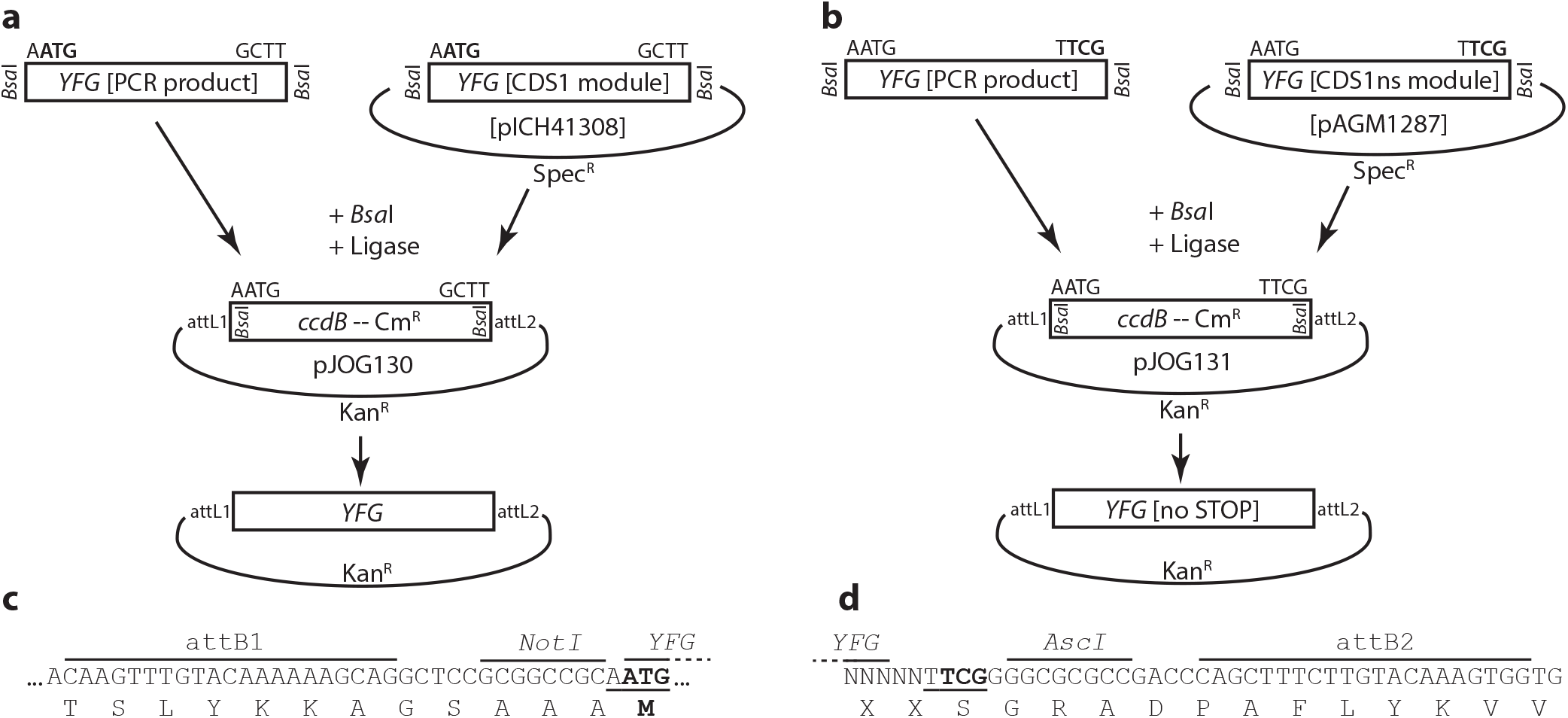
Generation of Gateway entry vectors by Golden Gate cloning. (a) Scheme of entry clone generation in pJOG130. Either PCR products flanked by BsaI restriction sites and suitable 4 bp overhangs or CDS1 Level 0 modules of the Modular Cloning system may be cloned into pJOG130 by BsaI cut/ligation in exchange for a *ccdB* cassette. (b) as in (a), but when using vector pJ0G131 for PCR products with suitable adaptors or CDS1ns modules of the Modular Cloning system. (c) Amino acid sequence encoded by att1 sites / adaptor sequences in pJOG130. The sequence created from using a pJOG130 derivative in a LR recombination reaction is shown. Translation will either initiate at an upstream START codon of an N-terminal epitope tag, or at the ATG codon depicted in bold if no N-terminal tag is fused during LR recombination. (d) as in (c), but when using a pJ0G131 derivative during LR recombination. Amino acid sequences encoded at 5’ fusion sites (att1) are equivalent as in (c) upon fusion of an N-terminal tag. The 3’ fusion site and respective linker sequences are shown. Sequences preceding the TCG (Ser) triplet depicted in bold, which is part of the Golden Gate overhang, will depend on design of PCR product or Level 0 module.

As mentioned, pJOG130/131 vectors are suitable for either toggling inserts from the Modular Cloning system to the Gateway system, or for creating Gateway entry clones from PCR products. In the first case, derivatives of pICH41308 (CDS1) or pAGM1287 (CDS1ns) containing inserts without internal *Bsa*I sites are used together with pJOG130 or pJ0G131, respectively, in *Bsa*I cut/ligation reactions (Fig. 1; Engler et al., 2014). These reactions are highly efficient (> 90% correct clones), and background-free by presence of the *ccdB* negative selection cassette in the target vector. The efficiency when cloning PCR products depends on (i) size and quality of the PCR product, and (ii) the number of internal *Bsa*I sites. PCR amplicons without internal *Bsa*I sites are cloned by simple *Bsa*I cut/ligation, and efficiencies are > 80 % when using high quality PCR products. However, also low-abundance PCR products can be cloned with reasonable efficiencies (> 20 %). Amplicons with internal *Bsa*I sites are cloned into pJOG130/131 by *Bsa*I cut/ligation, followed by enzyme denaturation and another round of ligation (cut/ligation-ligation). Even with two internal *Bsa*I sites, cloning efficiencies from 20-80 % are regularly obtained when using high-quality PCR products. In rare cases, overhangs generated by *Bsa*I restriction at internal sites may match vector overhangs of pJOG130/131. Internal restriction sites may be eliminated by splitting the desired insert in several amplicons, as previously described (Engler et al., 2008), but it might be advisable to follow a different cloning strategy in these cases. We would also advise on alternative strategies for amplicons with three or more internal *Bsa*I sites, as low efficiencies are expected. Summarizing, next to shuttling inserts from Modular Cloning to Gateway cloning, pJOG130/131 are intended for generation of novel gateway entry clones from PCR amplicons containing ≤ 2 internal *Bsa*I sites with a simple, generalized and cost-efficient cloning strategy (< 1 € per reaction). pJOG130 was successfully used for cloning of ~ 60 cDNAs encoding candidate interactors obtained in a yeast three hybrid screen to avoid sequence domestication prior to further confirmation of interactions.

### Standardized assembly of gateway destination vectors by modular cloning

Most labs relying on the Gateway cloning strategy dispose of a rich collection of Gateway destination vectors, and many different vector series are available to the community (e.g. Nakagawa et al., 2007; Grefen et al., 2010; Schlucking et al., 2013). However, e.g. the integration of novel fluorophores with altered properties or specialized demands may eventually necessitate the generation of novel destination vectors, often by cumbersome and inefficient cloning strategies.

We therefore developed three different Golden Gate modules (pJ0G387/267/562) containing the Gateway cassette (attR1-cat/ccdB-attR2), which enable systematic assembly of Gateway destination vectors for virtually any application following the standardized Modular Cloning grammar (Fig. 2). In Modular Cloning, Level 0 modules, which encode standardized parts of transcription units such as promoter, 5’UTR, coding sequence, C-terminal tag or 5’UTR and transcriptional terminator, are combined to a transcriptional unit in a respective Level 1 recipient by *Bsa*1-mediated Golden Gate cloning (Weber et al., 2011; Engler et al., 2014). pJ0G387 replaces a CDS1 module in a Golden Gate assembly reaction, and is designed for the generation of destination vectors coding for fusions with an N-terminal tag (Fig. 2a). It should be noted that assemblies with pJ0G387, but without an N-terminal tag module (NT1), are not appropriate and will lead to a modified N-terminus in final expression constructs. Similarly, pJ0G267 replaces CDS1ns modules in Level 1 assemblies. Depending on modules combined in the assembly reaction, pJ0G267 can be used for the construction of destination vectors coding for fusions with a C-terminal tag (Fig. 2b) or with C- and N-terminal tags (Fig. 2c). Finally, pJOG562 carries overhangs to replace promoter, 5’UTR and a CDS1ns module in assembly reactions. Here, assembly yields gateway destination vectors without promoter, designed for recombination of fragments encompassing promoter and coding sequence from a respective entry clone by LR reaction.

**Figure 2:**
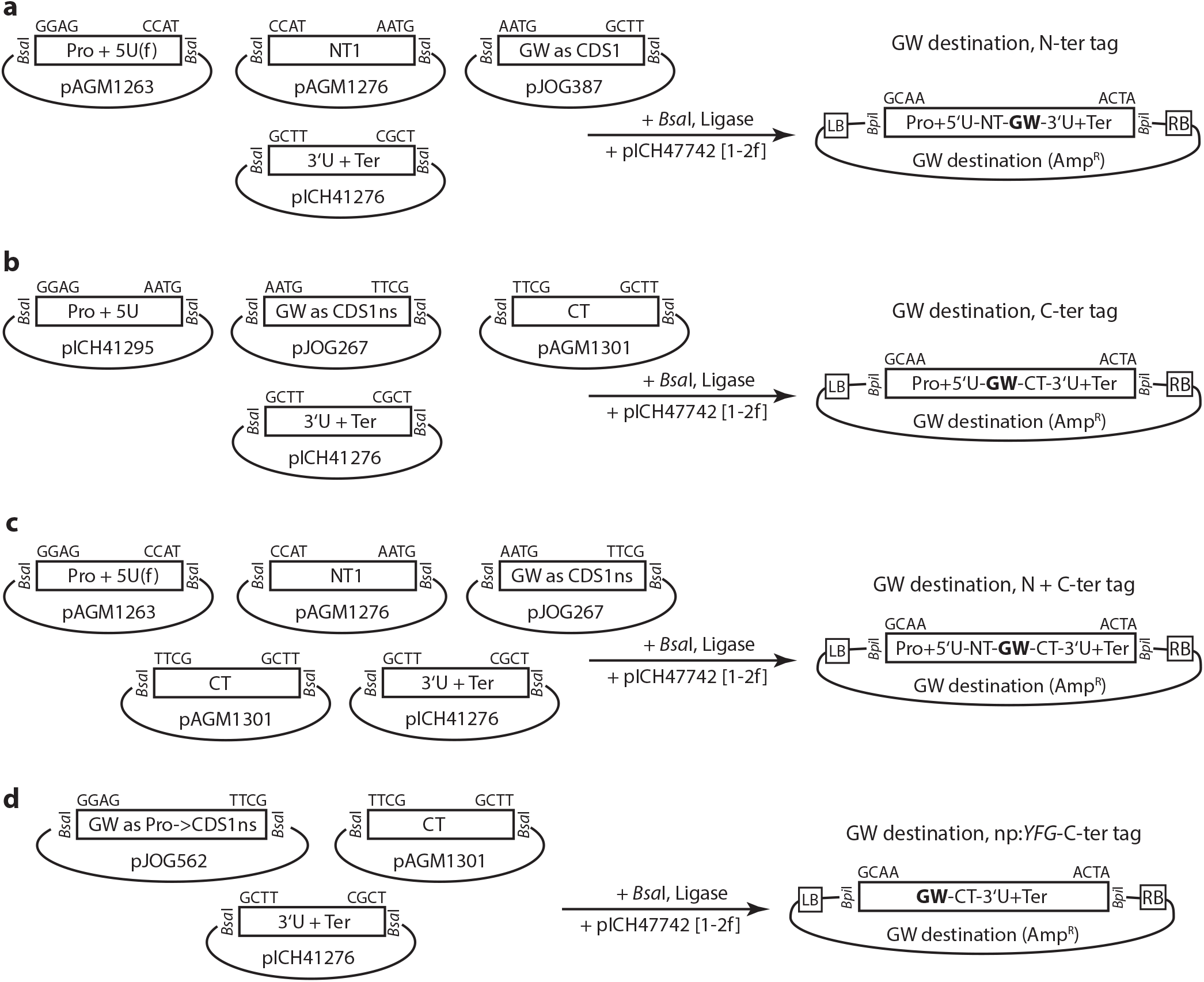
Assembly of simple Gateway destination vectors by Modular Cloning. (a) Assembly of Gateway destination vectors for N-terminal tagging of proteins of interest. (b) Assembly of Gateway destination vectors for C-terminal tagging of proteins of interest. (c) Assembly of Gateway destination vectors for N- and C-terminal tagging of proteins of interest. (d) Assembly of Gateway destination vectors for recombination of entry fragments containing both upstream regulatory sequences and a gene of interest, and for expression of C-terminally tagged proteins.

In the Modular Cloning system, Level 1 modules can be used for the generation of higher order Level 2 or Level M assemblies, in most cases to obtain multigene constructs (Weber et al., 2011; Engler et al., 2014). For hierarchical assemblies, transcriptional units can be assembled in 14 different Level 1 recipient vectors, and the used Level 1 recipient decides on orientation and position of a respective transcriptional unit in a final multigene construct. However, also Level 1 vectors are *Agrobacterium-*compatible, and assembled transcriptional units will be flanked by left and right borders (Fig. 2; Weber et al., 2011). Thus, Level 1 vectors can readily be used for e.g. *Agrobacterium*-mediated transient expression (“Agroinfiltration”), but do not contain a plant-selectable marker for stable transformation. For later combination with a plant-selectable marker, a Level 1 recipient for position 2 in a multigene construct in a forward orientation (1-2f) was used for the assembly of transcriptional units containing the Gateway cassette (Fig. 2). Plant-selectable marker cassettes were assembled as Level 1-1f modules (position 1, forward). In the following, a pnos:Bar-tnos cassette (BASTA selection) generated by standard Golden Gate assembly (see methods) was used, and several additional marker cassettes are available in the Modular Cloning format (Engler et al., 2014).

In a particular application, gateway destination vectors for fusions with a red fluorophore were required. Available destination vectors for fusions with DsRed did not produce satisfying results. New destination vectors containing the improved mCherry fluorophore were therefore assembled (Shaner et al., 2004). The Gateway modules pJOG387 and 562 were used in different Level 1 assemblies together with mCherry-coding modules. The resulting, Gateway cassette-containing transcriptional units (1-2f) were subsequently combined with a plant-selectable marker (1-1f) and a suitable end-linker in a Level M recipient (Fig. 3a), placing the plant-selectable marker next to the left border. Level 1 and level M assembly reactions were highly efficient; all tested clones were positive. The resulting plasmids were used for LR reactions, final expression vectors transformed into *Agrobacterium* and proteins transiently expressed in *Nicotiana benthamiana (Nbenth).* The newly assembled vectors were functional and facilitated reliable live cell imaging of calmodulin2 and IQ67 domain (IQD)8, in the cytosol and nucleus, and at the plasma membrane and microtubule cytoskeleton, respectively (Figs. 3b and c; Bürstenbinder et al., 2017). Conclusively, this standardized assembly scheme allows rapid generation of Gateway destination vectors from phytobricks.

**Figure 3:**
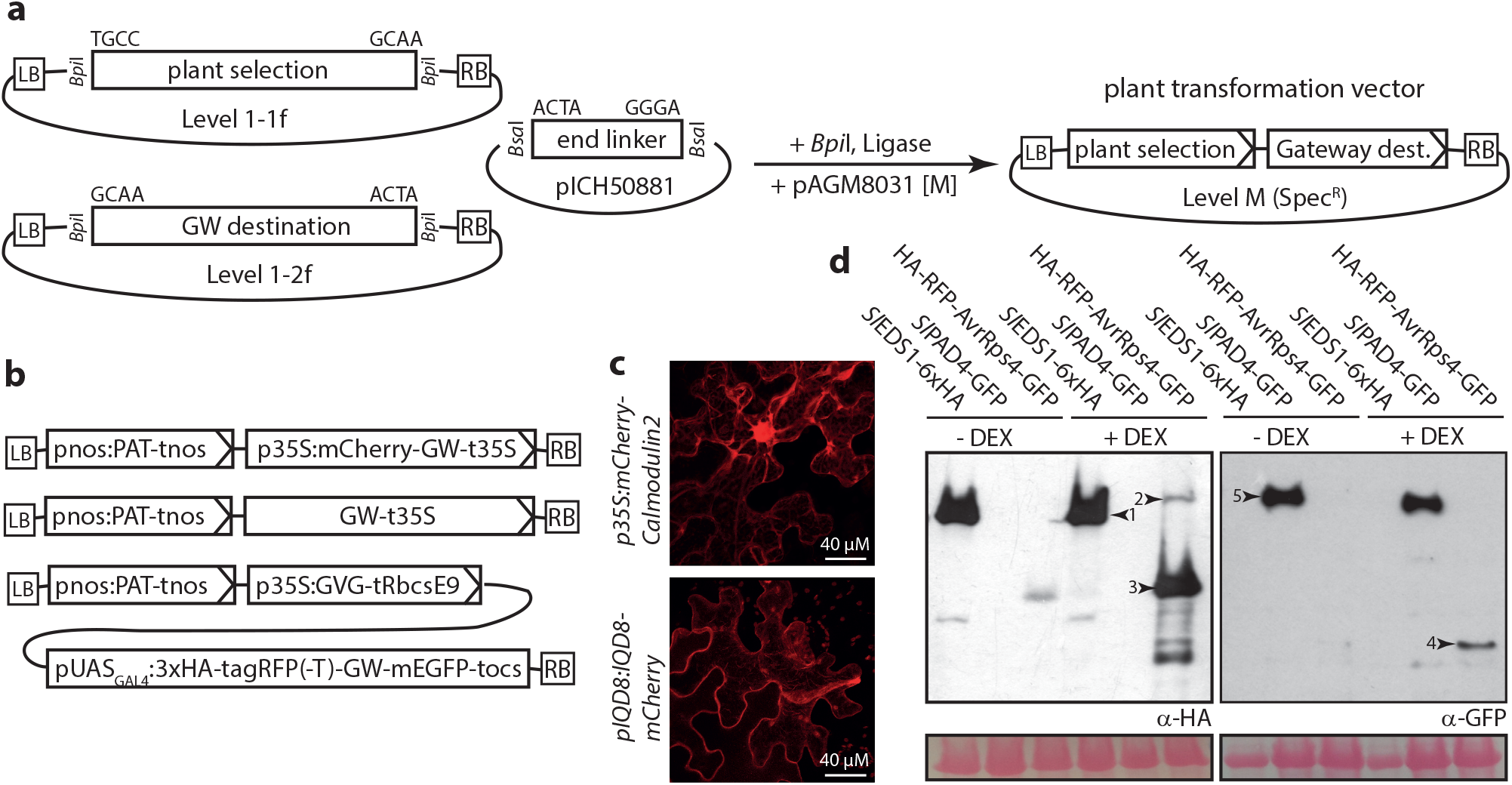
Assembly of Gateway plant transformation vectors by Modular Cloning and functional verification. (a) Exemplary scheme for assembly of simple plant transformation vectors containing a plant-selectable marker and a Gateway destination cassette. (b) Bi- and tri-partite plant transformation vectors assembled for functional verification. (c) Live-cell imaging of proteins transiently expressed in *N. benthamiana* from bi-partite transformation vectors shown in (b). Arabidopsis Calmodulin2 and IQD8 were expressed as fusions to mCherry, as indicated. Infiltrated leaf sections were analyzed 3 dpi. Maximum intensity projections are shown. (d) Immunoblot detection of proteins for functional verification of inducible expression vector shown in (b). Bands corresponding to the expected sizes of EDS1-HA (1), unprocessed HA-RFP-AvrRps4-GFP (2), processed HA-RFP-AvrRps4^N^ (3) and AvrRps4^C^-GFP and PAD4-GFP (5) are marked by arrowheads.

### Standardized assembly of gateway destination vectors II: Multipartite destination vectors

For a different application, a Gateway destination vector for dexamethasone (DEX)-inducible expression of a protein of interest with an N-terminal 3xHA-tagRFP(-T) (Shaner et al., 2008) and a C-terminal GFP tag was required. Plant transformation vectors for DEX-inducible expression consist of a plant-selectable marker, a fusion of an artificial transcription factor with the hormone-binding domain of the rat glucocorticoid receptor (“activator”), and a third transcriptional unit with the gene of interest under control of a synthetic promoter (“response element”). In presence of DEX, the artificial transcription factor is released from binding to cytosolic Hsp90, enters the plant cell nucleus, binds to the synthetic promoter and activates transcription of the response element (Picard, 1993). To our knowledge, the pTA7001 vector or derivatives are most commonly used for DEX-inducible *in planta* expression (Aoyama and Chua, 1997; McNellis et al., 1998). The modification of pTA7001 to our needs appeared impractical. We therefore decided to modularize components of the inducible system: GVG (GAL4-VP16-GR; activator) and pUAS (6xUAS_GAL4_-p35Smin; synthetic promoter). A transcription unit coding for the activator was assembled as Level 1-2f module, and a module for the response element containing the Gateway cassette was assembled as 1-3f module. Transcription units were assembled together with a plant-selectable marker and a suitable end-linker in a Level M recipient (Fig. 4a). We tested our new inducible expression vector by transiently expressing AvrRps4 from *Pseudomonas syringae* pv. *pisi* in *Nbenth.* (Hinsch and Staskawicz, 1996). AvrRps4 is cleaved *in planta* by a yet unknown plant protease (Sohn et al., 2009), which should lead to release of N- and C-terminal fragments of the 3xHA-tagRFP(-T)-AvrRps4-GFP fusion protein. Tomato EDS1 and PAD4 tagged with 6xHA and mEGFP, respectively, were expressed as control proteins, and detected both in presence and absence of DEX (Fig. 4d). AvrRps4 was strongly induced in presence of DEX, and N- and C-terminal fragments were detected by respective antibodies, confirming functionality of the newly constructed inducible expression vector (Fig. 4d). Thus, the presented strategy allows the assembly of simple or multipartite gateway destination vectors in a highly efficient manner and following the standardized Modular Cloning grammar.

**Figure 4:**
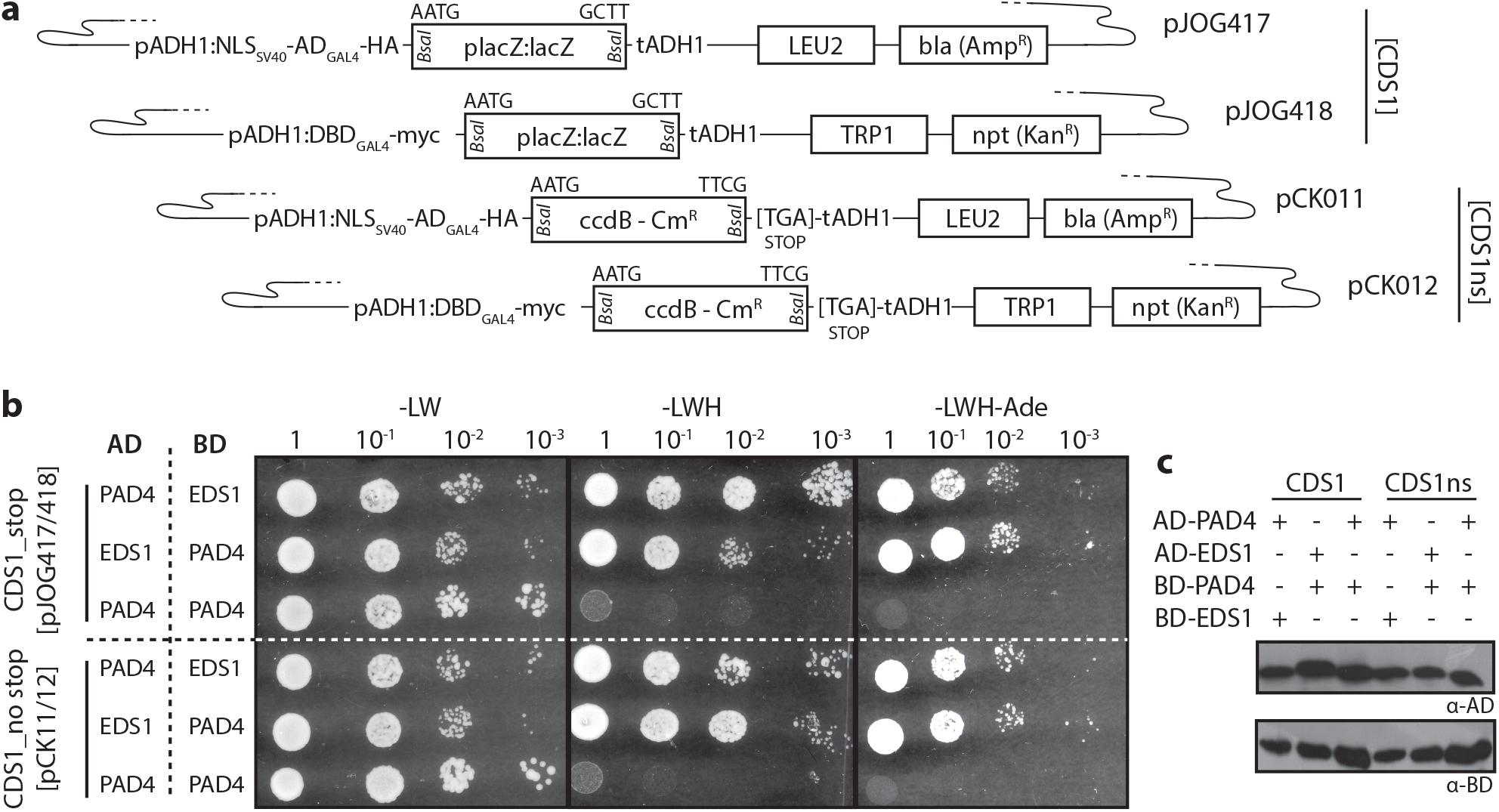
Modular Cloning-compatible vectors for a GAL4-based yeast two hybrid system. (a) Schematic depiction of yeast two hybrid vectors with most important features. (b) Functional verification of vectors shown in (a). Tomato *EDS1* and *PAD4* were mobilized into vectors shown in (a), resulting constructs co-transformed into yeast cells in the indicated combinations, and co-transformants grown in dilution series on media lacking leucine and tryptophan (-LW), or additionally lacking histidine (-LWH) and adenine (-LWH-Ade). (c) Immunoblot-detection of fusion proteins expressed in yeast cells in (b).

### Yeast Two Hybrid Vectors for Use with Level 0 Modules of the Modular Cloning Standard

The popular pGAD and pGBK vectors (Clontech), which are intended for GAL4-based yeast two hybrid interaction assays, were converted to the Modular Cloning standard (Fig. 4a). Bait and prey vectors were generated to receive CDS1 modules (pJ0G417/418) or CDS1ns modules (pCK011/012). The Y2H vectors for CDS1ns modules allow the use of identical Level 0 modules e.g. for *in planta* expression with a C-terminal epitope tag and for Y2H assays with minimal C-terminal modifications of yeast fusion proteins. A STOP codon directly follows the 3’ golden gate cloning overhang used for CDS1ns modules (T|TCG). Thus, yeast fusions proteins will contain as few as 1-2 additional amino acids depending on the design of the respective Level 0 module, and will terminate with a serine residue encoded by the TCG within the overhang. All vectors may also be used for Golden Gate cloning of PCR products carrying *Bsa*l adapters.

Vectors were tested using EDS1 and PAD4 from tomato. Arabidopsis EDS1 and PAD4 strongly interact to form a heterodimeric complex (Wagner et al., 2013). We had previously confirmed that tomato EDS1 and PAD4 also interacted in Y2H using Gateway-compatible pGAD/pGBK derivatives. CDS1 and CDS1ns Level 0 modules of tomato EDS1 and PAD4 were used for *Bsa*l cut/ligation reactions with the Y2H vectors, and all tested clones were positive in restriction digests. Resulting constructs were cotransformed into yeast, and primary transformants replica-plated on reporter media in dilution series (Fig. 4b). All yeast strains grew on –LW media selecting for presence of both plasmids in cotransformants. Growth on –LWH and –LWH-Ade media, indicative of interaction of bait and prey proteins, was observed upon co-expression of AD/BD fusions of EDS1 and PAD4 in either orientation, but not if PAD4 was tested for self-interaction (Fig. 4b), as previously observed. All fusion proteins were detected by immunoblotting (Fig. 4c), confirming integrity of the vector-encoded epitope tags and thus full functionality of the presented Y2H vectors.

### Bacterial type III secretion vectors for the modular cloning standard

Plant pathogenic bacteria often rely on the secretion of proteins directly into the cytoplasm of host cells via a type III secretion system (Büttner, 2016). Substrates for type III secretion are recognized by a yet enigmatic N-terminal secretion signal, and proteins normally not recognized as type III secretion substrates may be secreted via the type III system if fused to a respective signal. This has been extensively used to analyze e.g. the function of oomycete effectors in the “effector detector system” (Sohn et al., 2007; Fabro et al., 2011). Four different vectors for bacterial type III secretion and compatible with the modular cloning system were generated (Fig. 5a). Vectors contain either amino acids 1-134 of AvrRps4 and are thus very similar to the previously described pEDV vectors (Sohn et al., 2007), or amino acids 1-100 of the AvrRpt2 effector (Innes et al., 1993). With each of these secretion signals, a vector for CDS1 modules and for CDS1ns modules was generated, and ligation of CDS1ns modules results in a C-terminal 3xmyc epitope in final fusion proteins. The *Xanthomonas euvesicatoria* genes encoding the AvrBs3 and XopQ effectors were mobilized into CDS1 and CDS1ns vectors, respectively, for functional verification. AvrBs3 is a Transcription Activator-Like Effector (TALE), and AvrBs3-mediated induction of the *Bs3* resistance gene provokes a strong and rapid cell death reaction (Romer et al., 2007; Boch and Bonas, 2010). XopQ is recognized in the non-host plant *Nbenth,* and induces a mild cell death reaction (Adlung et al., 2016), which is abolished on an *eds1a-1* mutant *Nbenth* line (Ordon et al., 2017). Derivatives of the bacterial secretion vectors containing AvrBs3 or XopQ were mobilized into a *Pseudomonas fluorescence* strain carrying a chromosomal integration of the type III secretion system from *Pseudomonas syringae* (“EtHAn”; Thomas et al., 2009). Resulting strains were infiltrated into wild type, *Bs3* transgenic, and *eds1a-1* mutant *Nbenth* plants (Fig. 5b). AvrBs3-expressing strains provoked strong cell death on *Bs3* transgenic plants, as expected. This confirmed that both the AvrRps4- and AvrRpt2-derived secretion signals were functional. Similarly, XopQ-expressing strains provoked cell death reactions on wild type and *Bs3* plants, but not on *eds1a-1* plants (Fig. 5b). Notably, cell death reactions upon infiltration of AvrRpt2-XopQ strains were substantially and reproducibly stronger than those of AvrRps4-XopQ strains (Figs. 5b and S2), suggesting that either the AvrRpt2 signal might confer higher levels of protein translocation or the respective fusion protein might be more stable or more active. This demonstrates the utility of testing several different signals for bacterial translocation of a protein of interest into plant cells.

**Figure 5:**
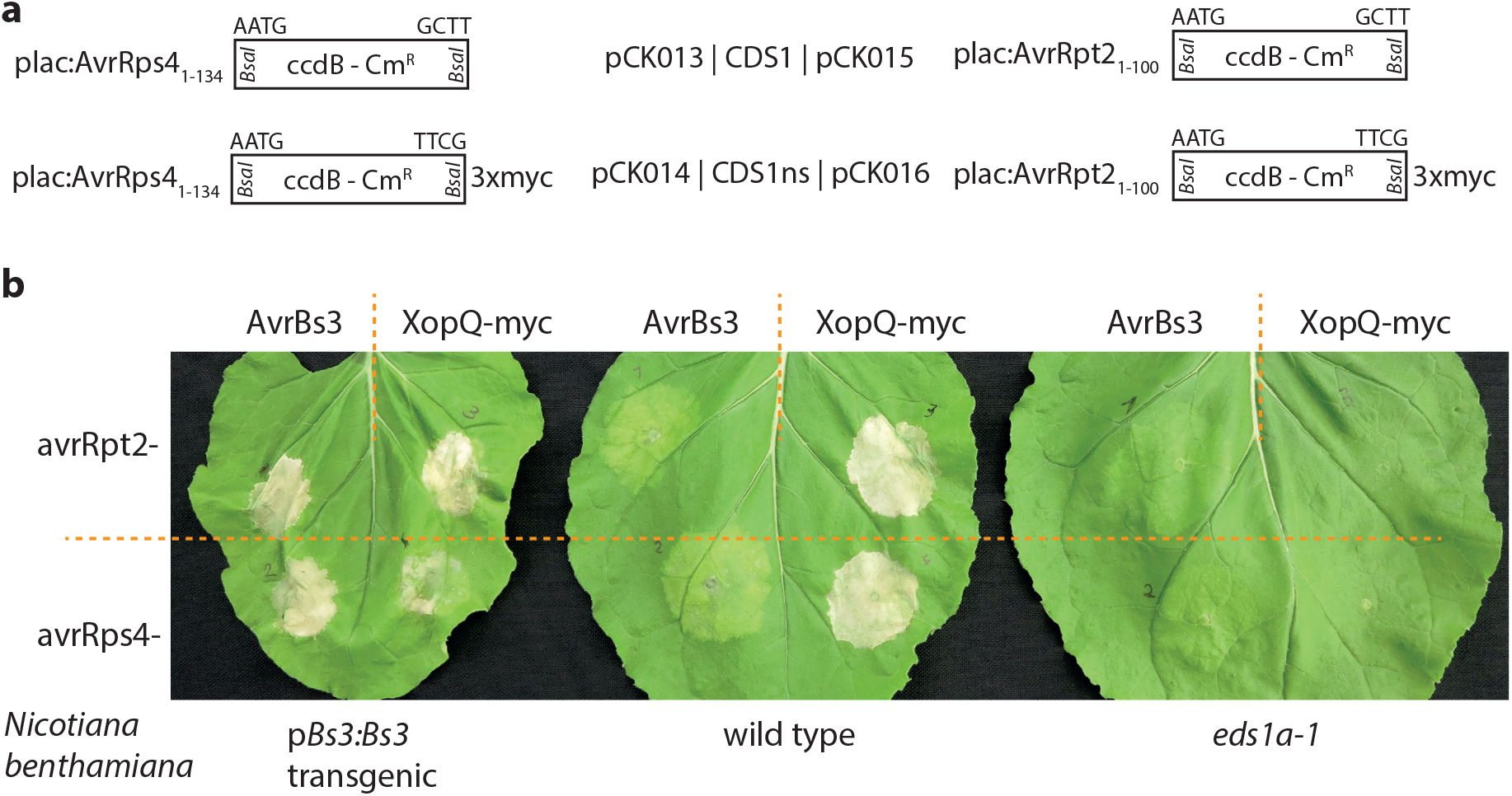
Bacterial type lll-delivery of proteins into *N. benthamiana* cells. (a) Schematic depiction of vectors for type lll-delivery of proteins. (b) Hypersensitive response induction assays for functional verification of vectors shown in (a). Either AvrBs3 or XopQ were cloned in vectors shown in (a), as indicated. Resulting constructs were mobilized into *Pseudomonas fluorescens,* strains inoculated at an OD_600_=0.4 on indicated *N. benthamiana* genotypes and symptoms documented 3 dpi.

An AvrRps4_1-136_ protein fragment was previously used to mediate bacterial translocation of cargo proteins, and fusions were at least partly processed in plants due to AvrRps4 cleavage by a plant protease (Sohn et al., 2007). *In planta* processing of proteins expressed from AvrRps4_1-134_ fusion vectors presented here was not tested. Irrespective of whether *in planta* processing occurs or not, final proteins will carry non-native N-termini and might also lack e.g. post-translational modifications, potentially impairing protein functions. Thus, we do not consider the likelihood for functionality of delivered proteins to increase through cleavage of secretion signals. Indeed, high cell death-inducing activity of AvrRpt2-XopQ, for which no cleavage is expected, demonstrates that secretion signals may have minimal and different effects on cargo functionality. To possibly avoid negative effects of the fused translocation signal on the delivered cargo moiety, the two fragments are fused by a Gly-Gly-Ser linker in pCK13-16 vectors presented here.

*Agrobacterium-mediated* expression of XopQ in wild type *Nbenth* plants induces mild chlorosis to mild necrosis, although the protein accumulates to high levels in plant tissues (Adlung et al., 2016; Schultink et al., 2017). In contrast, bacterial translocation of XopQ here induced a strong cell death response (Fig. 4b). Although not tested in detail, it is generally assumed that protein levels inside the plant cell obtained by *Agrobacterium-mediated* expression largely exceed those of bacterial translocation. Increased abundance of XopQ inside plant cells upon bacterial secretion is thus not a likely explanation for the phenotypic differences. As an alternative to protein dosage, we propose that a negative effect of *Agrobacterium* strain GV3101 on HR development (Li et al., 2017) or other peripheral constraints during transient, *Agrobacterium*-based assays (Erickson et al., 2014) might be at the basis of the observed differences in XopQ-induced HR development.

### A golden gate-cloning vector for Tobacco Rattle Virus-induced gene silencing

Virus-induced gene silencing is an attractive and convenient method for the rapid knock-down of a gene of interest without the need for transformation or gene knockout. A tobacco rattle virus-based system is most commonly used, and functional in a number of different plant species including tomato and *Nbenth* (Ratcliff et al., 2001; Liu et al., 2002). A fragment of the gene of interest is inserted into the RNA2 of the bipartite genome of TRV, and viral RNAs are reconstituted in the plant by expression from *Agrobacterium*-delivered T-DNAs. To facilitate rapid and cost-efficient cloning into a TRV RNA2 vector, an existing vector system (Liu et al., 2002) was adapted to Golden Gate cloning, and the cloning site was replaced by a **Bsa*I*-excised *ccdB* cassette as negative selection marker (Fig. 6a). For functional verification, a fragment of the *Nbenth PDS* gene was inserted into the previously used, gateway-compatible TRV2 vector and the newly generated Golden Gate-compatible vector. *PDS* encodes for Phytoene Desaturase essential for the production of carotenoids, and knockdown induces strong photo-bleaching of leaves. *Agrobacterium* strains carrying the respective TRV2 vectors were side-by-side co-inoculated with TRV1-containing strains into the lower leaves of *Nbenth* plants, and leaf bleaching observed 14 days later (Fig. 6b). Both TRV vectors induced leaf bleaching to similar extents, confirming functionality of the Golden Gate-compatible derivative. The pTRV2-GG vector was also used for silencing of the *Nbenth EDS1* gene, and the XopQ-induced HR was consistently abolished on *EDS1* knock-down plants in several independent biological replicates (Fig. 6c).

**Figure 6:**
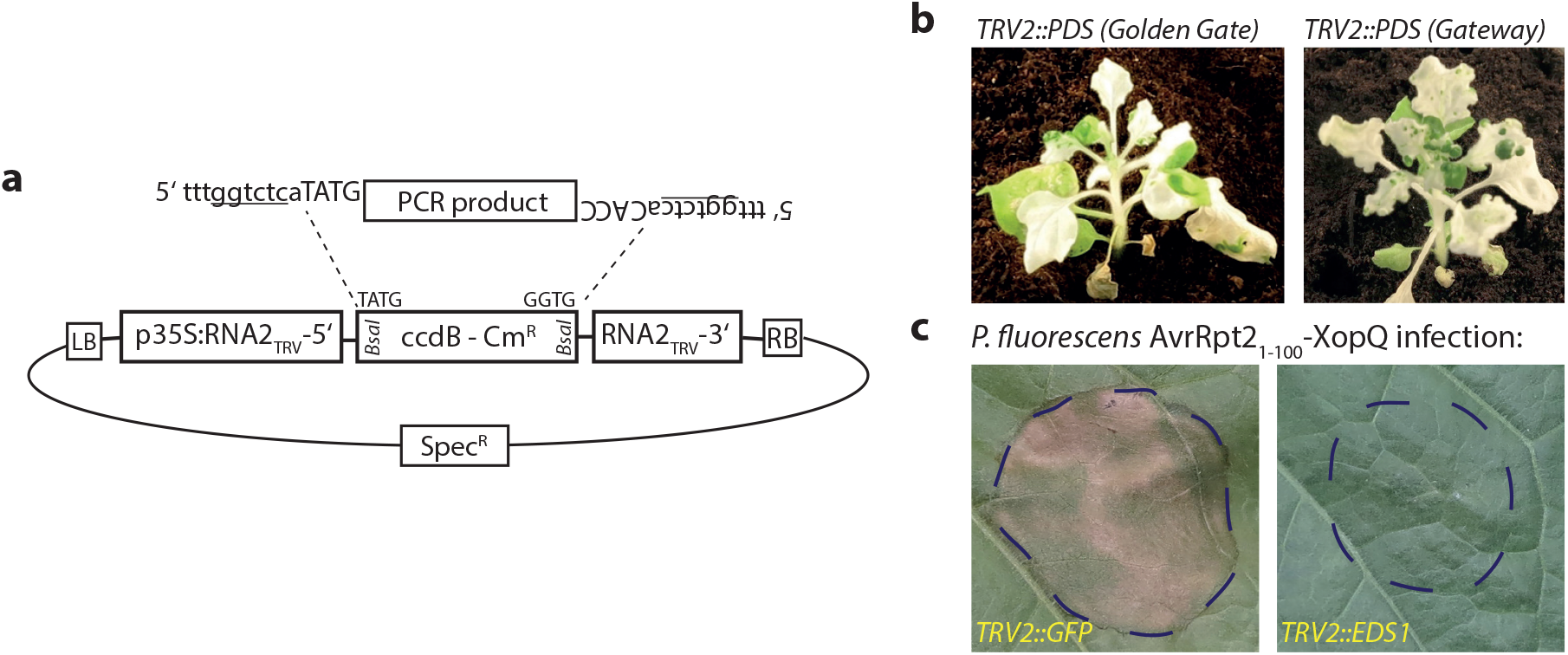
A Golden Gate cloning-compatible TRV2 vector for virus induced gene silencing. (a) Schematic depiction of pTRV2-GG. The adaptors required for introduction of PCR products are indicated. (b) Functional verification of pTRV2-GG by silencing of *PDS.* pTRV2 and a commonly used Gateway-compatible TRV2 vector containing identical *PDS* fragments were compared for silencing efficiencies. (c) Silencing of *NbEDSl* using a pTRV2-GG derivative. *Pseudomonas fluorescence* bacteria expressing an AvrRpt2_1-100_-XopQ fusion protein (OD_600_ = 0.2) were inoculated 14 days after inoculation of pTRV strains, and plant reactions were documented 3 dpi.

All vectors presented in this and previous sections are summarized in supplemental table S1. Annotated sequence files are provided in Appendix S1.

### An extended set of plant parts, or phytobricks, for the modular cloning system

Level 0 modules, or phyotobricks, are the building blocks and thus the limiting component for assemblies following the Modular Cloning grammar. As an extension to the previously released Plant Parts (Engler et al., 2014), we here provide ~ 80 additional Level 0 modules, summarized in supplemental table S2. These modules were experimentally verified as part of our ongoing projects if not indicated otherwise (Table S2), and functional data is presented for a few selected modules. The provided new phytobricks comprise a variety of module types, e.g. modules for inducible gene expression (Fig. 3), promoters for constitutive and tissue-specific gene expression in Arabidopsis (Fig. S3), transactivation (Fig. S4), additional fluorophores for (co-) localization and FRET analyses (Hecker et al., 2015), signals for modifying subcellular localization, or epitope tags. In addition to the Modular Cloning and Plant Parts toolkits, these modules will further enhance the versatility of this hierarchical DNA assembly system and facilitate its implementation in the plant research community. All vectors described (Tables S1 and S2) will be distributed via Addgene, and annotated nucleotide sequences (GenBank format) are contained in Appendix S1.

## Conclusions

Novel Golden Gate-based hierarchical cloning strategies, such as Modular Cloning, allow the rapid and cost-efficient assembly of simple transcriptional units or multigene constructs from basic building blocks (phytobricks). The underlying assembly standard, or molecular grammar, ensures efficient bioengineering by re-utilization and sharing of phytobricks. Accordingly, ~ 80 novel phytobricks are provided to foster this idea of shared resources. Furthermore, we show how the Modular Cloning assembly standard may, by integrating just a few modules, also be used for inexpensive generation of Gateway entry clones, toggling between cloning systems, or standardized assembly of Gateway destination vectors. These alternative applications of Modular Cloning may be particularly helpful to avoid the eventually laborious domestication of sequences at early stages of a project, as e.g. a first screening of candidate genes, or to connect resources available for different cloning systems.

One major advantage of Gateway cloning consists in the availability of destination vectors for virtually any biological system or experimental setup. In contrast, Modular Cloning and GoldenBraid were so far mainly designated for the generation of plant expression/transformation constructs. Similar hierarchical DNA assembly systems were also developed for e.g. yeast or prokaryotes (Lee et al., 2015; Moore et al., 2016), but only some rely on the same fusion sites between building blocks for assembly (Perez-Gonzalez et al., 2017). Here, we present vectors for direct use of Modular Cloning Level 0 modules in yeast interaction assays or for bacterial translocation into plant cells. Similarly, new vectors need to be adapted to this cloning standard in the future, e.g. for protein production in *Escherichia coli.* This will ensure seamless and efficient integration of synthetic biology standards and novel DNA assembly strategies, and will streamline laboratory workflows by reducing molecular cloning workloads.

## Material and Methods

### Plant material, growth conditions, bacterial infection assays and virus induced gene silencing

*Nicotiana benthamiana* wildtype, *eds1a-1* mutant plants (Ordon et al., 2017) and *pBs3:Bs3* transgenic plants (Schreiber et al., 2015) were cultivated in a greenhouse with 16 h light period, 60% relative humidity at 24/20°C (day/night). For transient *Agrobacterium-mediated* expression, plate-grown bacteria were resuspended in Agrobacterium infiltration medium (AIM; 10 mM MES pH 5.7, 10 mM MgCl_2_) to an OD_600_=0.4 or as indicated, and infiltrated with a needleless syringe. For imaging of IQD8 and Calmodulin2, *Agrobacterium* strains were mixed in a 1:1 ratio with a strain for expression of p19. Plasmids were mobilized into a *Pseudomonas fluorescens* strain containing a chromosomally-encoded *Pseudomonas syringae* type III secretion system (“EtHAn”; Thomas et al., 2009) by triparental mating, and plate-grown bacteria were resuspended in 10 mM MgCl2 prior to infiltration.

For virus induced gene silencing, *Agrobacterium* solutions were infiltrated in the bottom leaves of three week-old plants. Photo-bleaching was documented 14 d later, or plants were used for challenge inoculations.

### Yeast two hybrid assays, immunoblotting and live cell imaging

Derivatives of pGAD and pGBK vectors (pJ0G417-418 and pCK011-pCK012) were co-transformed into frozen competent yeast cells of strain PJ69-4a as previously described (Gietz and Schiestl, 2007). Single colonies were cultivated in liquid SD media for 48 h, and dilution series prepared. Yeast cell solutions were plated on selective media using a multichannel pipette, and plates were grown for 3-4 days prior to documentation. For immunoblot detection from yeast, proteins were extracted as previously described (Kushnirov, 2000). For extraction of plant proteins, leaf discs were ground in Laemmli buffer and boiled at 92 °C for 5 minutes. Proteins were separated on SDS-PAGE gels, transferred to nitrocellulose membranes, and detected via HRP-conjugated secondary antibodies (GE Healthcare) using Supersignal West Pico and Femto substrates (Pierce). Primary antibodies used were α-GFP (mouse monoclonal), α-HA (rat monoclonal; both from Roche and now distributed by Sigma), α-AD (GAL4 activation domain) and α-BD (GAL4 DNA-binding domain; both mouse monoclonal; Takara). Imaging was performed either on a Zeiss LSM 700 inverted microscope using a 40x water immersion objective, or a Zeiss LSM780 system. For imaging of IQD8 and Calmodulin2, mCherry was excited with a 555 nm laser, and emission was detected between 560 and 620 nm. Images are maximum intensity projections of z stacks. For simultaneous imaging of mTRQ, mEGFP and mCherry, fluorophores were excited with 458, 488 and 561 nm lasers, and emission was detected between 463-482, 499-543 and 587-630nm. For simultaneous imaging of mEGFP and chlorophyll A, 488 and 633 nm lasers were used for excitation, and emission was detected between 490-517 and 656-682 nm.

### Molecular cloning

Vectors for generation of gateway entry vectors (pJOG130-131) were generated by ligating a PCR amplicon encoding for a *ccdB* cassette and flanked by *Bsa*l sites cutting respective 4 bp overhangs into the *Ascl/Notl* sites of a pENTR/D derivative. Gateway modules (pJ0G267, 387) were generated by ligation of an attR1-ccdB/cat-attR2 PCR amplicon into pAGM1287 and pICH41308 (Engler et al., 2014), respectively, or into the *EcoRV* site of a custom cloning vector (pJ0G397) for generation of pJ0G562. For generation of GAL4-based yeast two hybrid vectors (pJ0G417-418), *Bsa*l sites in the backbones of pGAD and pGBK vectors (Clontech) were eliminated by mutagenesis, and a lacZ cassette was subsequently ligated into the *EcoRI/XhoI* sites. The pCK011-12 vectors were derived from these by replacing the lacZ cassette by a *ccdB* cassette with respective adaptors. The bacterial secretion vectors are based on a Golden Gate-compatible pBRM derivative (Szczesny et al., 2010), and secretion signals and *ccdB* cassette were ligated into the *BsaI/EcoRI* sites. To generate pRNA2-GG, the 5’ and 3’ fragments of TRV2 and a *ccdB* cassette were cloned between 35S promoter and terminator sequences in pVM_BGW (Schulze et al., 2012). All Level 0 modules (Table S2) were constructed as described (Engler et al., 2014), and internal restriction sites eliminated. For ligations or Golden Gate reactions, generally 20 fmole of all components were used, and reactions performed as previously described (Weber et al., 2011). Additional details and primer sequences are available upon request.

---

**Supplemental Figure S1:**
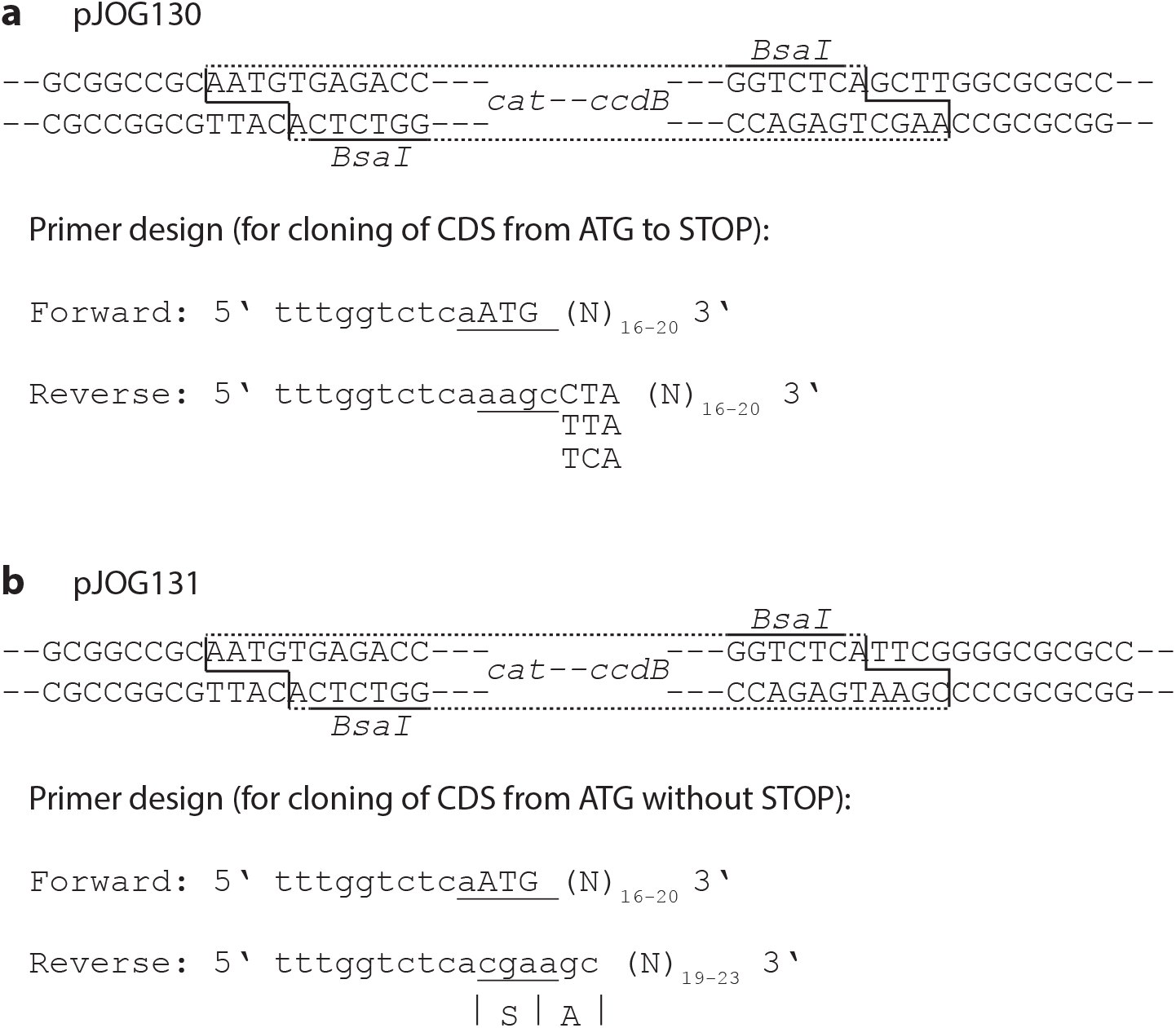
Primer design for cloning into Golden Gate-compatible entry vectors pJOG130/131. (a) The *ccdB* cassette contained in pJOG130 with *Bsa*l restriction sites underlined is shown. The adaptors required for PCR amplification of suitable fragments are depicted below. Underlined sequences represent the 4 bp overhangs utilized for Golden Gate cloning, and Ns represent the gene specific portion of respective PCR primers. (b) as in (a), but for pJ0G131.

**Supplemental Figure S2:**
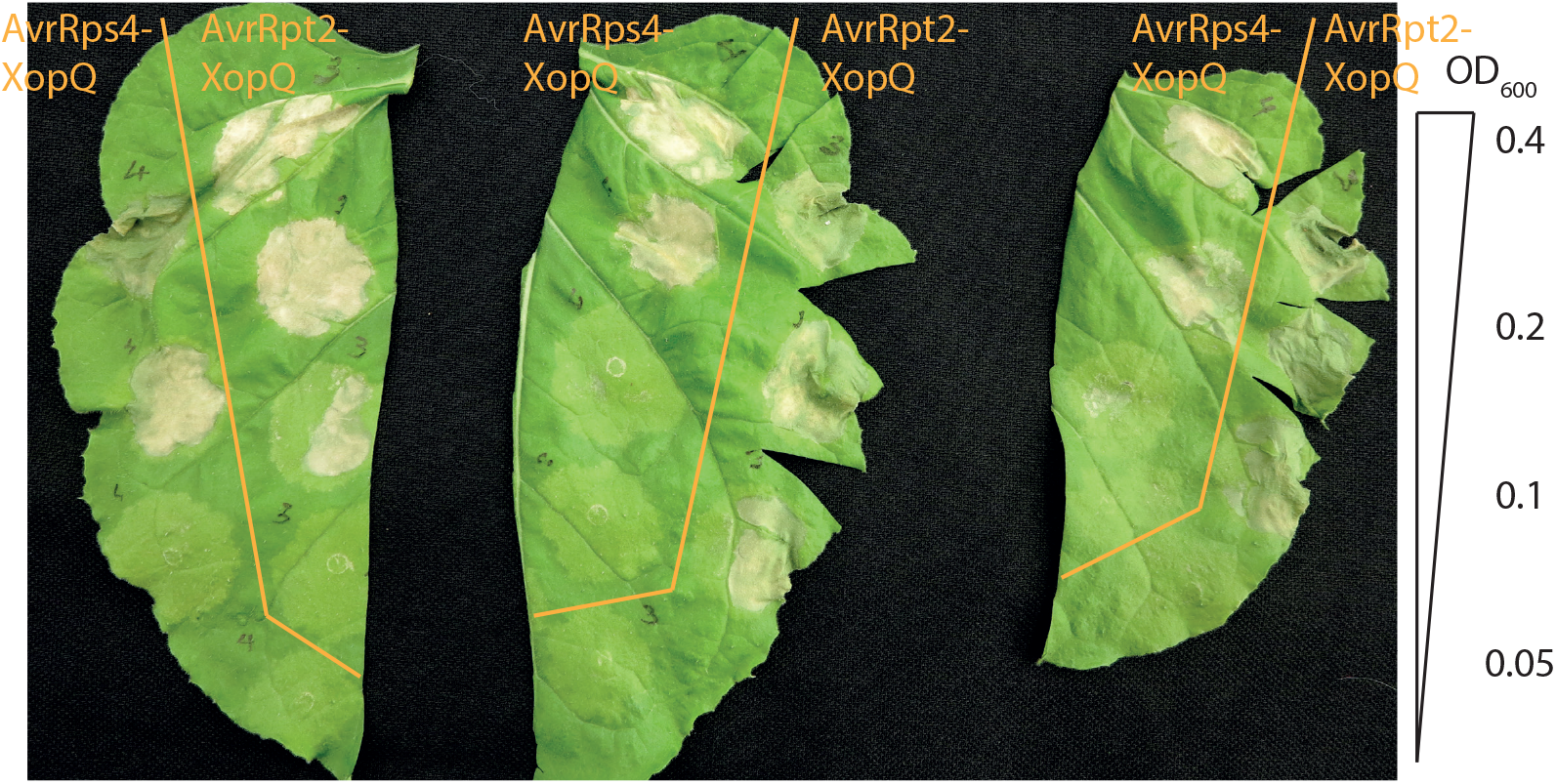
Enhanced hypersensitive response induction by AvrRpt2-XopQ fusions. *Pseudomonas fluorescens* strains translocating either AvrRpt2_1-100_-XopQ or AvrRps4_1-134_-XopQ fusions were inoculated into wild type *N. benthamiana* plants, and symptom formation was documented 3 dpi. Four different bacterial densities, ranging from OD_600_=0.4-0.05 were used, and were infiltrated descendingly in the indicated leaf sections.

**Supplemental Figure S3:**
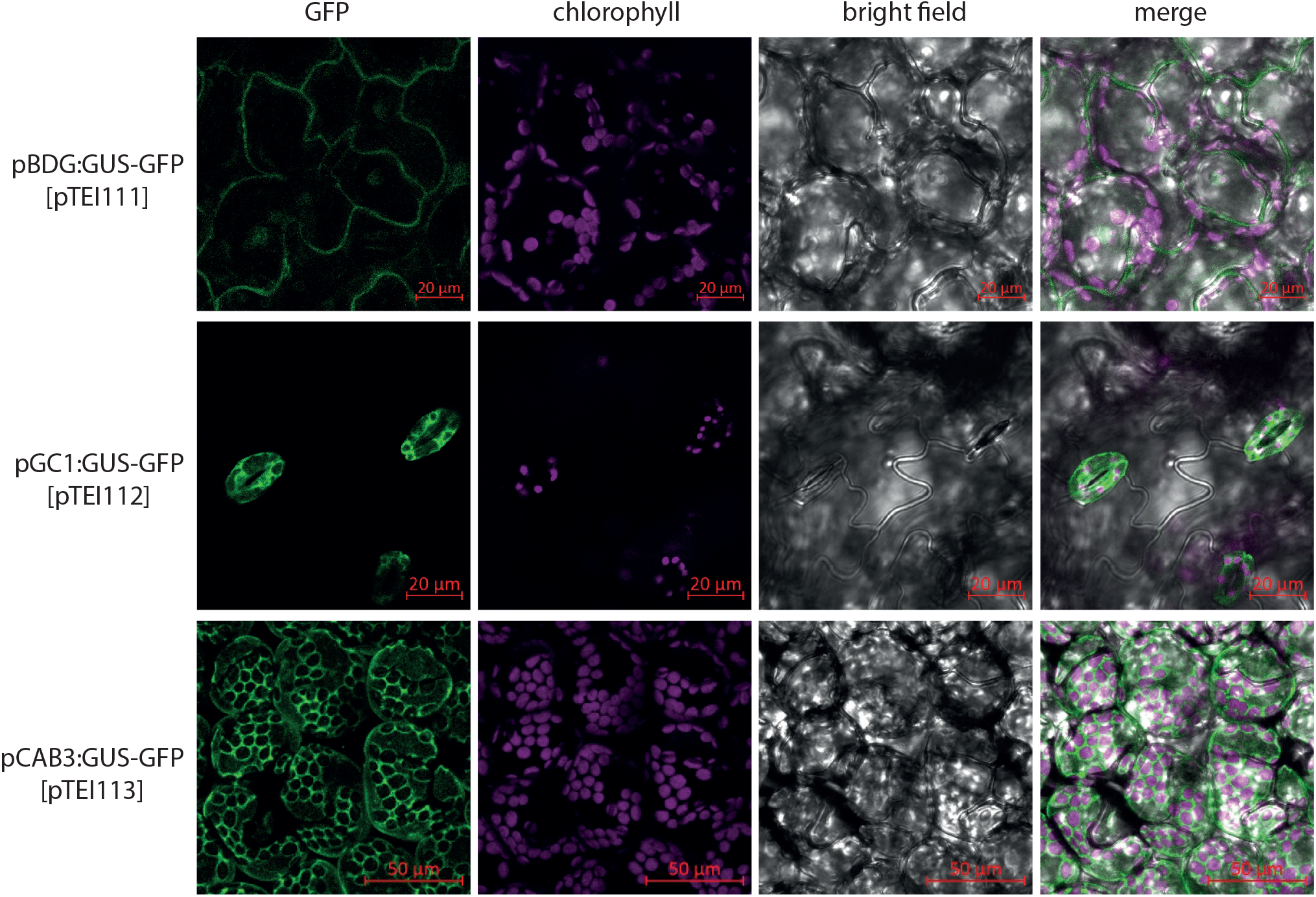
Promoter fragments for tissue-specific gene expression in Arabidopsis leaves. Transgenic Arabidopsis plants expressing GUS-GFP under control of the indicated promoter fragments were generated, and three-week-old T_1_ plants analyzed by confocal laser scanning microscopy. Maximum intensity projections of z-stacks are shown. Three independent T_1_ plants were analyzed for each construct with similar results.

**Supplemental Figure S4:**
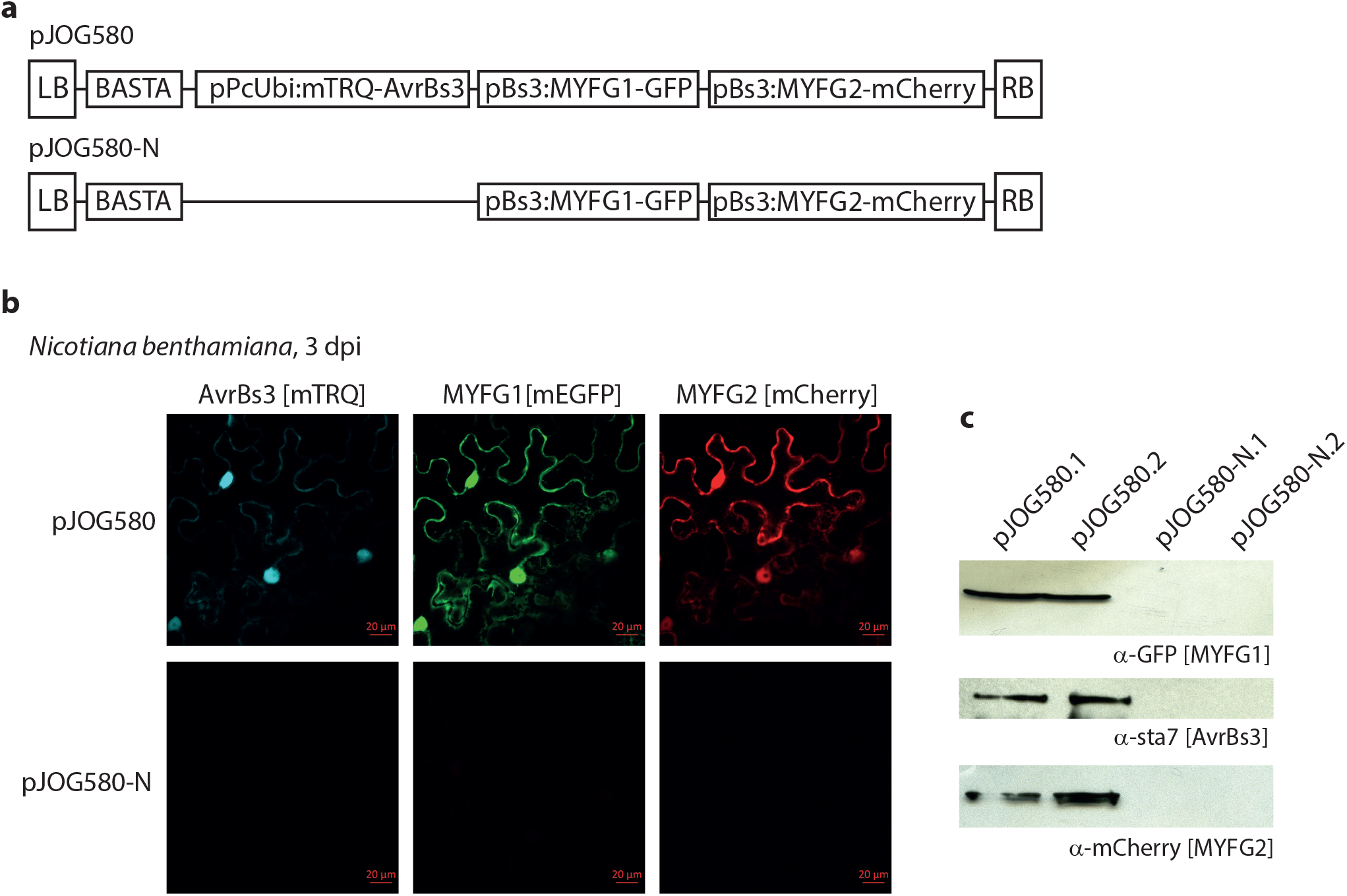
Utilization of TALEs for tightly regulated, high-level transactivation. (a) Schematic drawing of transactivation constructs used for transient expression. (b) Strong and specific transactivation of TALE-controlled genes. *Agrobacterium* strains containing constructs depicted in (a) were infiltrated into *N. benthamiana.* Leaf tissues were analyzed by confocal laser-scanning microscopy 3 dpi. (c) Immunoblot analysis of protein extracts prepared from leaf tissues analyzed in (b).

**Supplemental Table S1:**
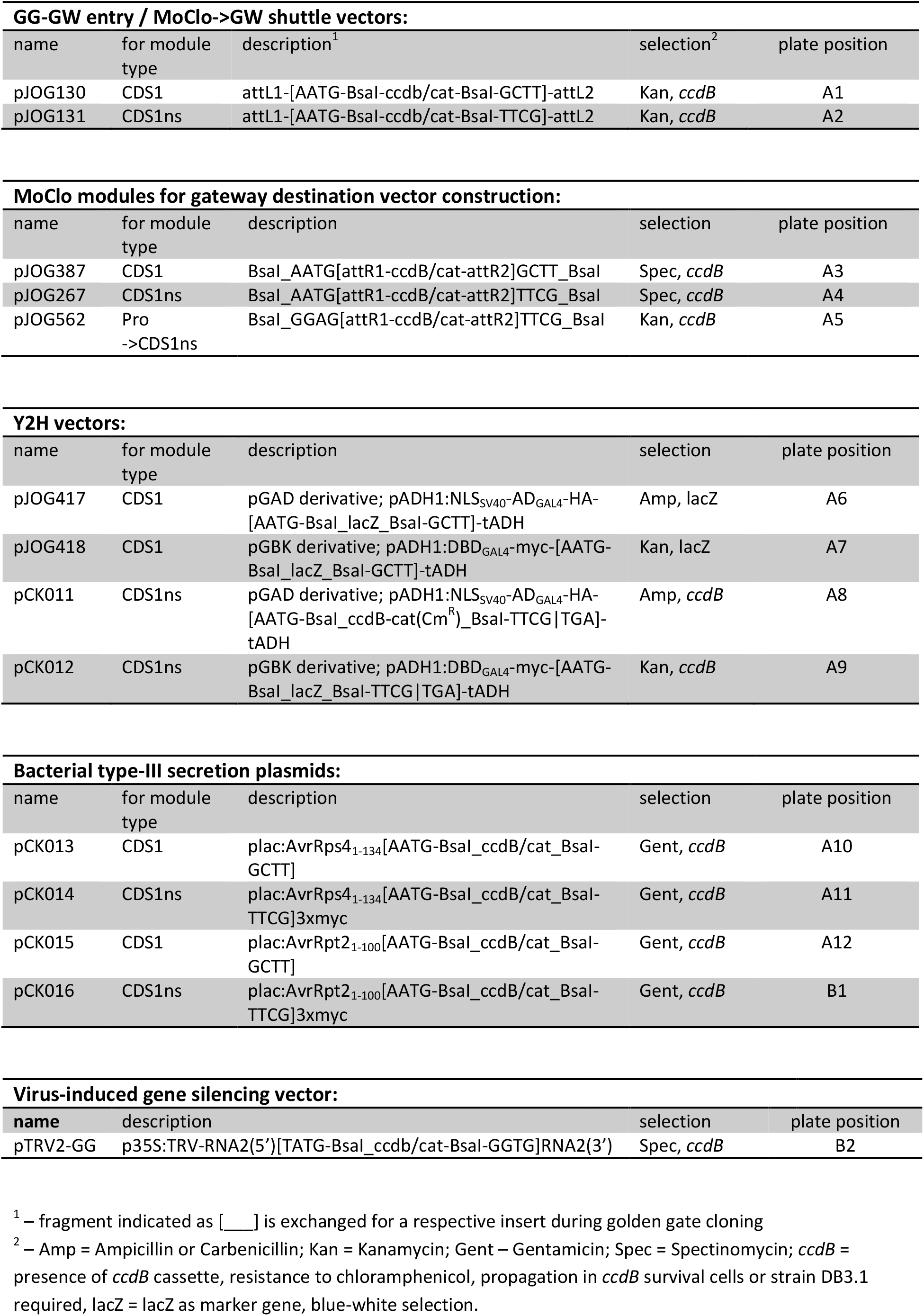
Modular cloning-compatible vectors for specialized applications

**Supplemental Table S2:**
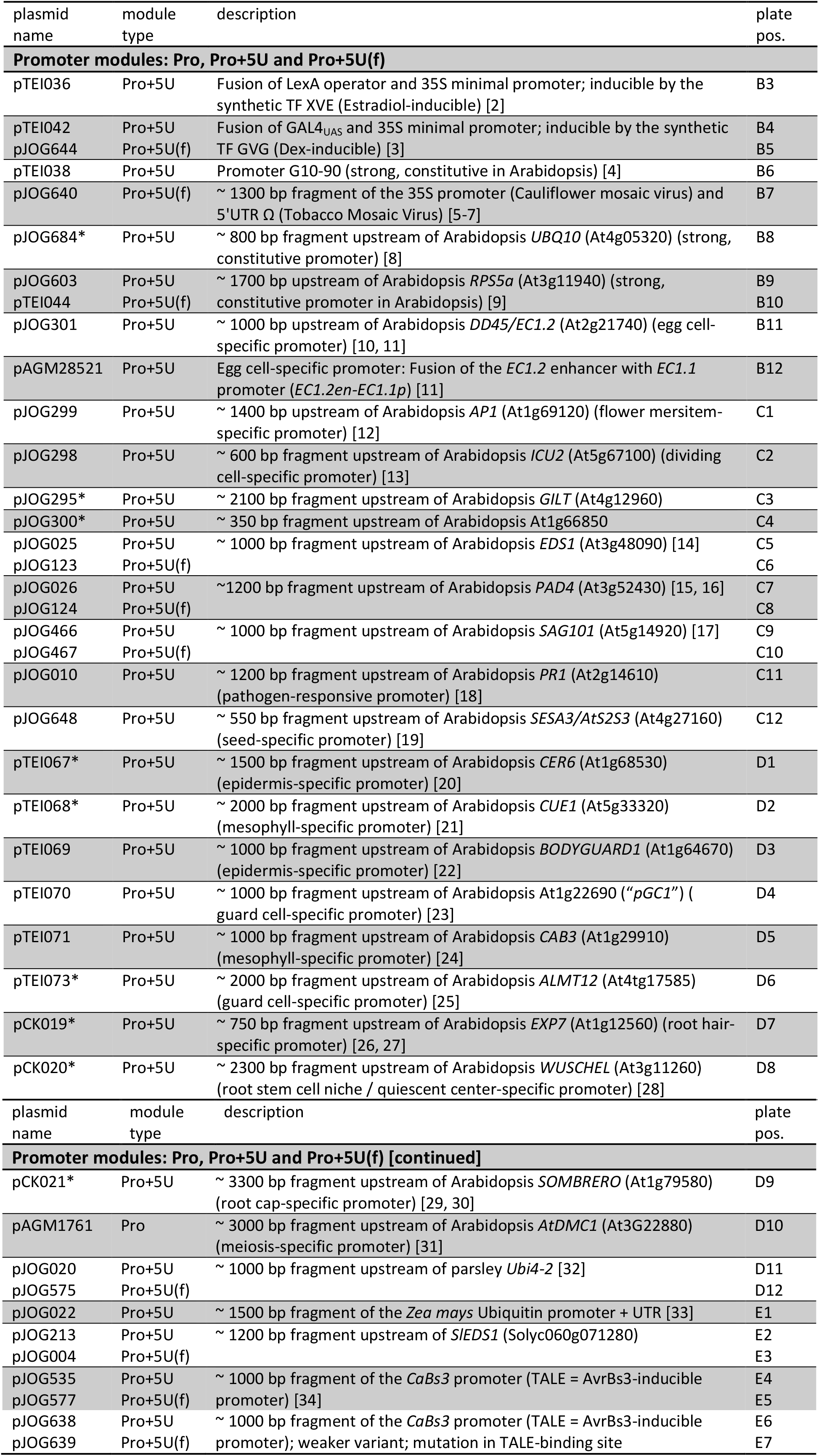
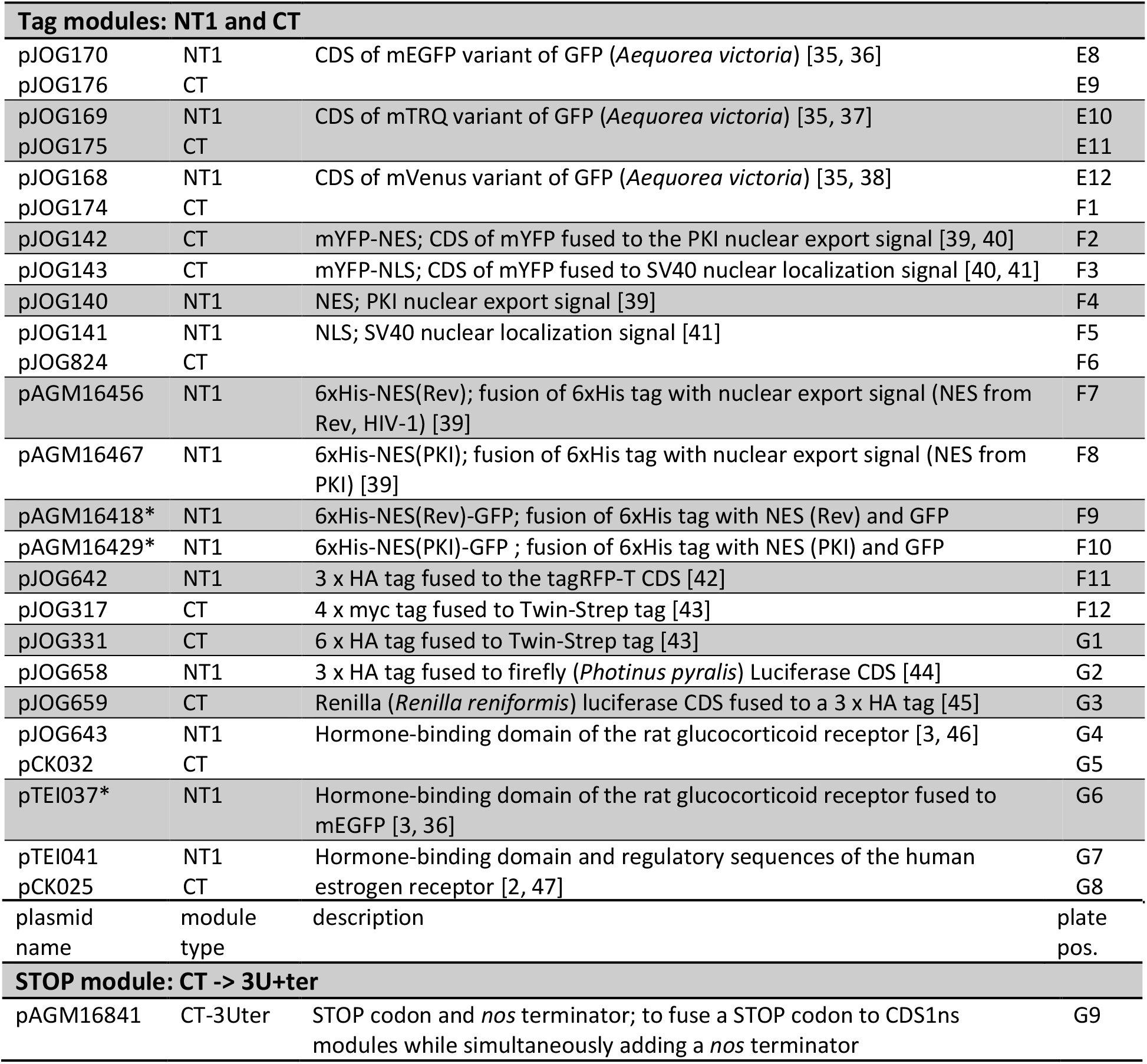
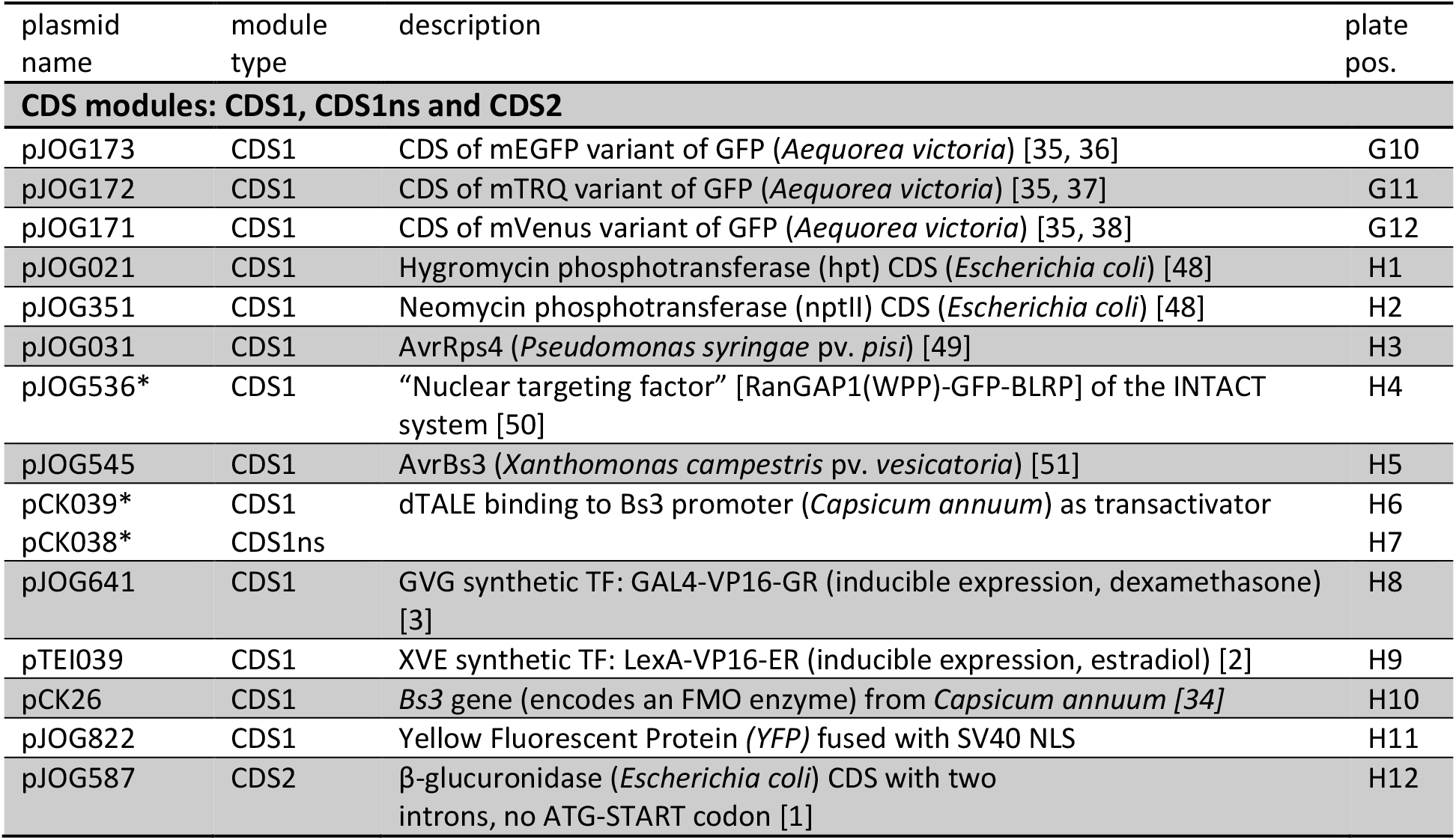
Modular cloning Level 0 modules

Supplemental Table S3: Promoter fragments for tissue-specific gene expression in Arabidopsis leaves

Appendix S1: Archive containing annotated sequence files (GenBank format) for all provided DNA modules.

## Author contributions

JG, JO, TI and JS planned and performed experiments and analyzed data. CK performed experiments. CL and YD provided pCK19-21 Arabidopsis promoter modules. SM and RG provided all pAGM*** modules. KB tested GW mCherry vectors. JS wrote the manuscript with contributions from all authors.

## Acknowledgements

We thank Christian Grefen for providing 2in1 FRET vectors, Annett Richter for providing a pZmUbi-containing plasmid, Tom Schreiber for providing AvrBs3 modules, Jens Boch for providing AvrRps4 and AvrRpt2-containing plasmids, all of which were used as PCR templates. Christine Wagner, Lennart Schwalgun and Samuel Grimm are acknowledged for technical assistance, and Jessica Erickson for assistance with CLSM imaging. We also thank Bianca Rosinsky foralways providing excellent plants. This work was funded in part by GRC grant STU 642-1/1 to Johannes Stuttmann.

